# An integrated single-cell RNA-seq atlas of the mouse hypothalamic paraventricular nucleus links transcriptional and functional types

**DOI:** 10.1101/2023.07.19.549666

**Authors:** J.B. Berkhout, D. Poormoghadam, C. Yi, A. Kalsbeek, O.C. Meijer, A. Mahfouz

## Abstract

The hypothalamic paraventricular nucleus (PVN) is a highly complex brain region that is crucial for homeostatic regulation through neuroendocrine signalling, outflow of the autonomic nervous system, and projections to other brain areas. The past years, single-cell datasets of the hypothalamus have contributed immensely to the current understanding of the diverse hypothalamic cellular composition. While the PVN has been adequately classified functionally, its molecular classification is currently still insufficient.

To address this, we created a detailed atlas of PVN transcriptional cell types by integrating various PVN single-cell datasets into a recently published hypothalamus single-cell transcriptome atlas. Furthermore, we functionally profiled transcriptional cell types, based on relevant literature, existing retrograde tracing data and existing single-cell data of a PVN-projection target region.

In our PVN atlas dataset, we identify the well-known different neuropeptide types, each composed of multiple novel subtypes. We identify *Avp*-*Tac1, Avp*-*Th, Oxt*-*Foxp1, Crh*-*Nr3c1* and *Trh*-*Nfib* as the most important neuroendocrine subtypes based on markers described in literature. To characterize the pre-autonomic functional population, we integrated a single-cell retrograde tracing study of spinally-projecting pre-autonomic neurons into our PVN atlas. We identify these (pre-sympathetic) neurons to co-cluster with the *Adarb2* ^+^ clusters in our dataset. Finally, we identify expression of receptors for *Crh, Oxt, Penk, Sst*, and *Trh* in the dorsal motor nucleus of the vagus, a key region that pre-parasympathetic PVN neurons project to.

Concluding, our study present a detailed overview of the transcriptional cell types of the murine PVN, and provides a first attempt to resolve functionality for the identified populations.

## 1. Introduction

The hypothalamus is a highly complex brain region that is crucial for homeostatic regulation. It consists of several nuclei that are functionally and molecularly diverse. Single-cell RNA sequencing has contributed immensely to the current understanding of the cellular composition of the hypothalamus. A recently published single-cell atlas based on 17 single-cell datasets is the current most high-resolution insight into the cellular makeup of the murine hypothalamus [27]. Yet, not all hypothalamic nuclei are represented at equal resolution in this atlas.

One of the underrepresented nuclei is the paraventricular nucleus (PVN), a small nucleus of the anterior hypothalamus, adjacent to the third ventricle. It is composed of several neuropeptidergic neuron types, mainly expressing arginine-vasopressin (AVP), oxytocin (OXT), corticotropin-releasing hormone (CRH), thyrotropin-releasing hormone (TRH) and somatostatin (SST) [4]. These are involved in various homeostatic processes, including the regulation of hormonal axes through neuroendocrine neurons, regulation of the ANS through pre-autonomic neurons, and regulation of – *inter alia* – behavior through centrally-projecting neurons [4, 8, 30, 16].

The neuroendocrine subset of neurons is usually described as being subdivided in the large magnocellular and smaller parvocellular neurons projecting to either the posterior pituitary or median eminence. The magnocellular neurons release the neurohormones AVP and OXT into the systemic bloodstream via the posterior pituitary, whereas parvocellular neurons release CRH, TRH and SST in the median eminence to be transported to the anterior pituitary via the hypophyseal portal system. In the anterior pituitary, CRH/AVP, TRH and SST act uon the different populations of trophic cells to stimulate or inhibit the hypothalamus-pituitary-adrenal (HPA), hypothalamus-pituitary-thyroid (HPT) and hypothalamus-pituitary-somatic (HPS) axes, respectively.

The pre-autonomic neurons project to the sympathetic and parasympathetic preganglionic neurons in the intermediolateral nucleus (IML) of the spinal cord and dorsal motor nucleus of the vagus (DMV) in the brainstem, respectively. In addition, these pre-autonomic neurons project to other brainstem nuclei such as the nucleus of the solitary tract, the central gray and the raphe nucleus [4]. Though existence of these projections has been described, the molecular identity of these neurons has not been completely elucidated yet. Several studies have attempted to uncover the molecular identity of the pre-autonomic neurons in the PVN, by utilizing retrograde tracing in combination with immunohistochemistry or in situ hybridization [4, 23, 12, 6]. These studies have revealed the expression of several neuropeptides, like AVP, OXT, enkephalin (PENK), and dynorphin (PDYN) in these neurons. However, these studies were only able to characterize roughly half of retrogradely traced neurons, thus necessitating further characterization of the neuropeptides and other markers involved.

Neuropeptidergic projections to other central locations upstream of the brainstem exist as well. While not all circuits have been described as extensively, it has been known for some time that – for instance – AVP and OXT are involved in modulation of social behaviors. Whereas AVP can promote aggressive behaviors [8], OXT promotes postpartum maternal behavior [30], and various other regions have been reported to be part of oxytocinergic circuits [11]. Recently, one of these social behavior-modulatory circuits was characterized in detail by Lewis *et al*. [16], describing a subset of PVN *Oxt* ^+^ neurons that mediate social reward learning through projections to the nucleus accumbens.

While the PVN has been adequately classified functionally (**Fig. 1A**), the molecular classification is currently insufficient. Even less well-described is the relation between the molecular and functional classifications. For instance, it is not known whether the centrally projecting neurons are a separate population of PVN neurons or a subset of the neuroendocrine and/or pre-autonomic neurons. This lack of insight in the complexity of this region is impeding research progress, since targeting a single functional pathway is difficult without knowledge of the transcriptional identity of the different functional types. To address this issue, we aimed to create a comprehensive atlas of PVN transcriptional types from existing scRNA-seq datasets. Furthermore, we aimed to associate transcriptional and functional types based on literature of neuroendocrine neuron-enriched genes, existing retrograde tracing data and a single-cell dataset of a PVN-projection target region.

**Figure 1:**
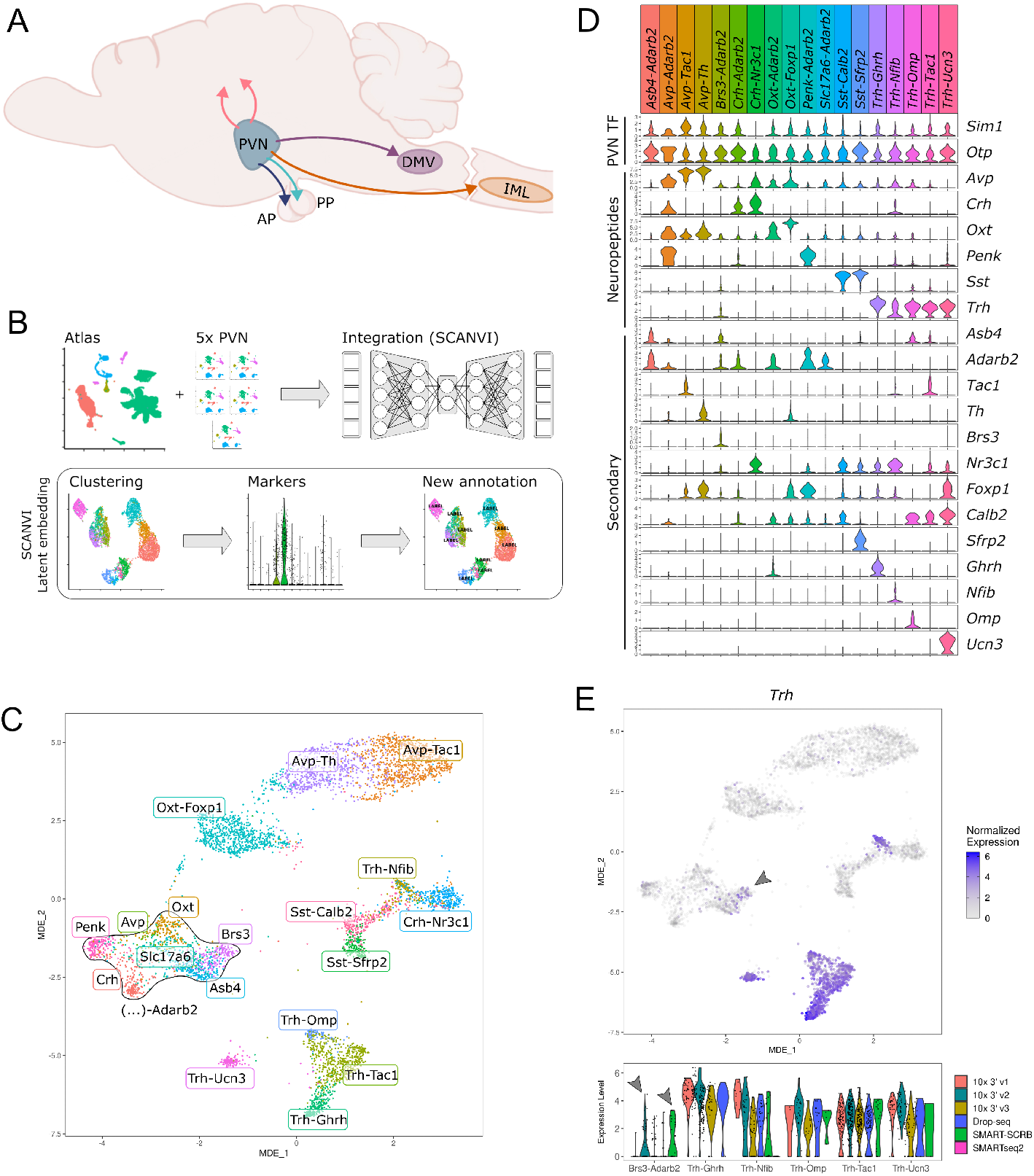
Neuropeptidergic PVN neurons consist of multiple subpopulations. **A**: Schematic showing the various projections from the PVN. Arrows point to the dorsal motor nucleus of the vagus (DMV), intermediolateral nucleus (IML), anterior pituitary (AP) and posterior pituitary (PP). Other (central) projections are denoted with the upward pink arrows. **B**: Schematic representation of the methods used for this section. Five PVN datasets were integrated with the hypothalamus atlas (HypoMap) using SCANVI. The integrated low-dimensional embedding was used for clustering and visualization. Clusters were annotated based on differential gene expression. **C**: Minimally distorting embedding (MDE) representation of the datasets, with the cluster annotation based on neuropeptide markers and secondary gene markers. **D**: Violin plots of the expression of PVN-specific transcription factors (TFs), neuropeptides and secondary cluster markers. **E**: MDE showing the expression of *Trh* in the dataset. Both point color and opacity are dependent on *Trh* expression to allow for identification of sparse populations. Shown below: violin plots of *Trh* expression in *Brs3* -*Adarb2* and all *Trh*^+^ populations, split by sequencing technique. Arrows indicate that *Trh* is expressed in *Brs3* -*Adarb2* cells, but in a sequencing method-dependent manner.

## 2. Results

### 2.1. PVN Neuropeptidergic neurons contain multiple subpopulations

To create the PVN atlas, we supplemented the whole-hypothalamus atlas [27] with various PVN and neuropeptide-specific single-cell datasets, that were originally not included in the atlas [16, 17, 25, 26, 33]. These datasets were integrated with SCANVI [32], and PVN neurons were selected from the integrated dataset. Finally, the resulting subset was clustered, and the identified clusters were annotated based on enriched genes (**Fig. 1B**).

After integration and annotation of the dataset, we first set out to characterize the neuronal populations and subpopulations. All expected major neuropeptidergic neuron populations were found (**Fig. 1C-D**), as well as some non-neuropeptidergic populations. Three populations in the dataset were excluded based on in-situ expression data: *Tcf7l2* -*Shox2, Slc32a1* -*Dlx1* and *Trh*-*Zic1* neurons (**Fig. S1A**). Anatomically, markers for these populations are found primarily in the thalamus and/or subparaventricular zone, especially compared to the rare immunoreactive neurons within the PVN (**Fig. S1B-D**). Thus, the relevance of these neurons for PVN function is unclear, given the low frequency of occurrence in the PVN, and their high abundance in bordering regions. All remaining clusters were positive for the vesicular glutamate transporter gene *Slc17a6*, and lacked the vesicular GABA transporter gene *Slc32a1* (**Fig. S2A-B**). Notably, despite the lack of *Slc32a1*, some of the neurons were positive for GABA biosynthesis enzymes, in particular for *Gad2* (**Fig. S2D**).

The identified clusters were annotated as neuropeptidergic subtypes, except the *Asb4* -*Adarb2, Brs3* - *Adarb2* and *Slc17a6* -*Adarb2* populations, which were not annotated as neuropeptidergic due to relatively low neuropeptide expression levels. Neuropeptidergic populations were further divided into subpopulations based on secondary markers. The *Avp*^+^ and *Oxt* ^+^ neurons were found to often present some level of coexpression of both genes (**Fig. 1D**), despite this being rarely observed at protein-level [9]. These clusters were characterized as either *Avp*^+^ or *Oxt* ^+^, based on which of the two genes was most abundantly expressed. This yielded *Avp*-*Adarb2, Avp*-*Tac1, Avp*-*Th, Oxt* -*Foxp1*, and *Oxt* -*Adarb2* as the subpopulations of *Avp*^+^ and *Oxt* ^+^ neurons (**Fig. 1D**). We further identified one population of *Penk* ^+^ neurons, *Penk* -*Adarb2*, two subpopulations of *Crh*^+^ neurons, *Crh*-*Nr3c1* and *Crh*-*Adarb2*, and two subpopulations of *Sst* ^+^ neurons, *Sst* -*Calb2, Sst* -*Sfrp2* (**Fig. 1D**).

Finally, the *Trh*^+^ neurons were split in multiple categories: *Trh*-*Ghrh, Trh*-*Nfib, Trh*-*Omp,Trh*-*Tac1*, and *Trh*-*Ucn3* (**Fig. 1D**). Notably, while the *Brs3* -*Adarb2* cluster was not annotated with a neuropeptide, it actually did express *Trh* (**Fig. 1E**). This expression was present at relatively low levels compared to the other *Trh*^+^ clusters and was found in a sequencing method-dependent manner (**Fig. 1E**). As such, the *Brs3* -*Adarb2* population was included in further analyses pertaining to *Trh*^+^ populations.

### 2.2. Neuroendocrine markers are enriched in distinct neuropeptide subpopulations

We next set out to identify which clusters may be linked to neuroendocrine function. In each neuropeptidergic type (**Fig. 2A**), genes previously identified to be expressed in neuroendocrine cells were analyzed for presence in subpopulations. Furthermore, we compared similarities between subpopulations, revealing that not all neurons within a neuropeptide class were transcriptionally most similar to each other (**Fig. 2B**). For instance, the *Oxt* -*Foxp1* neurons were more similar to the *Avp*^+^ neurons than the *Oxt* -*Adarb2*. In turn, the *Oxt* -*Adarb2* neurons were most similar to other *Adarb2* ^+^ neurons (**Fig. 2B**).

**Figure 2:**
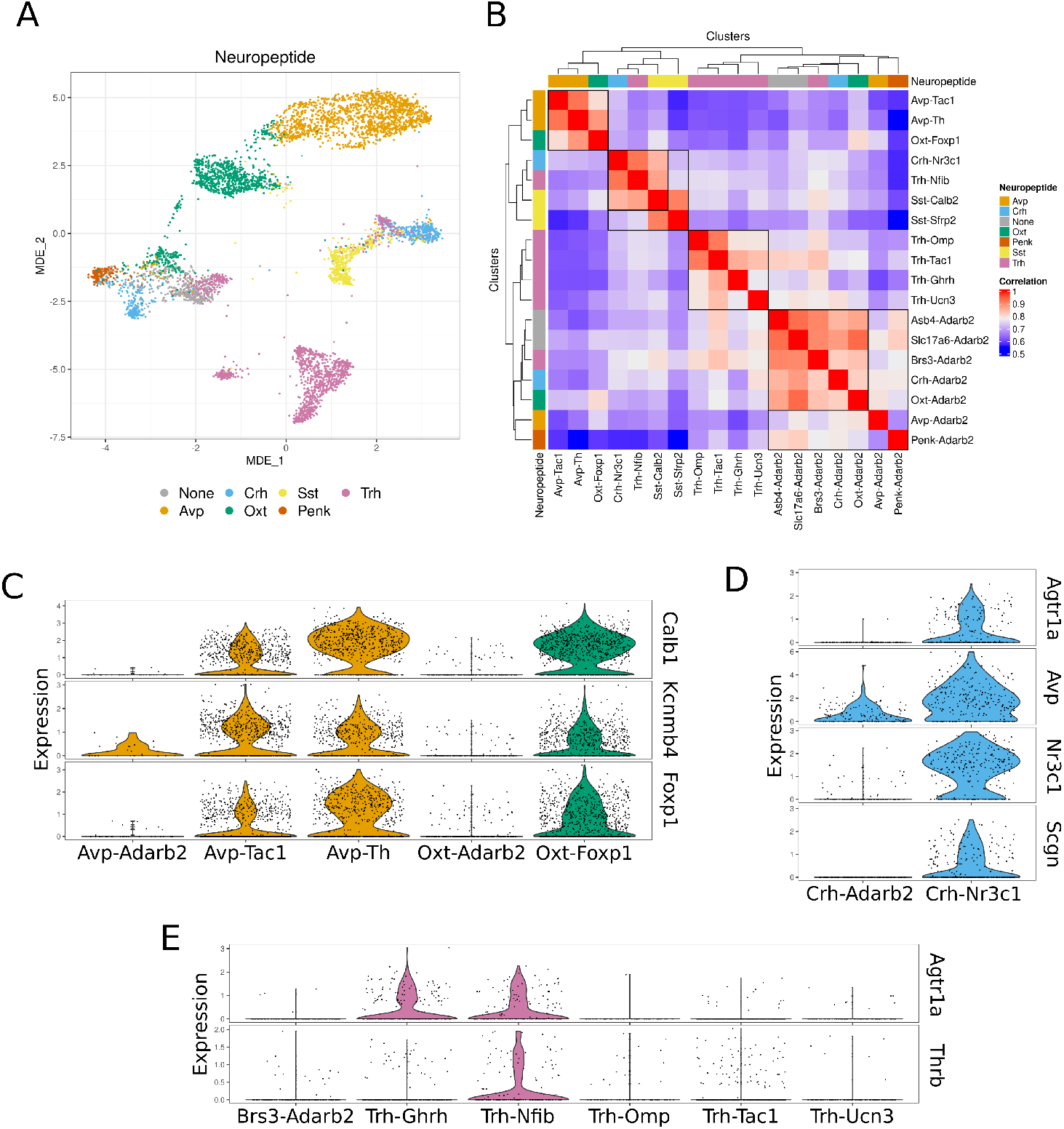
Expression of neuroendocrine markers in subpopulations of *Avp*^+^, *Oxt* ^+^, *Crh* ^+^ and *Trh* ^+^ neurons.. **A**: MDE representation of the data, colored by neuropeptide class. Coloring identical to violin plot colors in panels **(C)-(E) B**: Heatmap of correlations between clusters, calculated on pseudobulk expression values of highly variable genes. **C**: Violin plots of the expression of neuroendocrine *Avp*^+^ and *Oxt* ^+^ markers. **D**: Violin plots of the expression of neuroendocrine *Crh*^+^ markers. **E**: Violin plots of the expression of neuroendocrine *Trh*^+^ markers.

To functionally classify the *Oxt* ^+^ neurons, we used the neuroendocrine *Oxt* ^+^ markers as described by Lewis *et al*. [16]. In their study, a large overlap between magnocellular electrophysiology and neuroendocrine projection *Oxt* ^+^ neurons was found, and this magnocellular neuroendocrine population was reported to be enriched in *Calb1, Kcnmb4* and *Foxp1*. Notably, a closer inspection of their single-cell data reveals that their magnocellular neuroendocrine cluster contains both *Avp*^+^ and *Oxt* ^+^ neuron (**Fig. S3**). Thus, these markers for the magnocellular neuroendocrine population are relevant to not only *Oxt* ^+^ neurons, but also *Avp*^+^ neurons.

In our data, these magnocellular neuroendocrine markers were found to be enriched in the *Oxt* -*Foxp1* cells relative to the *Oxt* -*Adarb2* cells, suggesting the former to be the neuroendocrine population (**Fig. 2C**). Both *Avp*-*Tac1* and *Avp*-*Th* clusters were found to be strongly correlated with the magnocellular neuroendocrine *Oxt* ^+^ neurons (**Fig. 2B**), whereas the *Avp*-*Adarb2* cluster was not. Furthermore, the *Avp*-*Tac1* and *Avp*-*Th* populations were enriched in the magnocellular neuroendocrine markers relative to *Avp*-*Adarb2* (**Fig. 2C**). Between *Avp*-*Tac1* and *Avp*-*Th*, the *Avp*-*Th* population was slightly more enriched in these markers, but the relative differences were smaller than the difference to the *Avp*-*Adarb2* population or between *Oxt* ^+^ subpopulations.

Both the high degree of similarity between the *Avp*-*Tac1* and *Avp*-*Th* populations, as well as the expression of neuroendocrine markers in both populations imply *Avp*-*Tac1* and *Avp*-*Th* may be magnocellular neuroendocrine *Avp*^+^ neurons. It is currently not known if the differences between *Avp*-*Tac1* and *Avp*-*Th* neurons are functionally relevant, as the distinction has not been described before. It should be noted that, although positive for *Th*, the *Avp*-*Th* neurons do not express *Ddc* (**Fig. S4A**). Thus, these neurons cannot perform the final dopamine biosynthesis conversion, and therefore are not dopaminergic.

For the *Crh*^+^ neurons, multiple markers for neuroendocrine function have been described. First and foremost, the glucocorticoid receptor (*Nr3c1*; GR) is an essential mediator of regulatory feedback in *Crh*^+^ neurons of the HPA axis [14]. Another key component of these neurons is *Avp*, which strongly potentiates CRH-driven ACTH release from the anterior pituitary [10]. Further, *Scgn* has been proposed as a marker of neuroendocrine *Crh*^+^ cells, as *Scgn* silencing abolishes stress-driven ACTH release [21]. Lastly, *Agtr1a* has been found to be expressed in neuroendocrine *Crh*^+^ and *Trh*^+^ neurons [7]. All these markers were all enriched in the *Crh*-*Nr3c1* population, suggesting these neurons to be the neuroendocrine *Crh*^+^ population (**Fig. 2D**).

Like in *Crh*^+^ neurons, neuroendocrine *Trh*^+^ neurons are reported to express *Agtr1a* [7]. Additionally, literature indicates that the thyroid hormone receptor beta (THRB) is critical to the T_3_-mediated negative feedback on *Trh* expression [19]. Beside these two, markers for neuroendocrine *Trh*^+^ cells have not been described in literature. While several *Trh*^+^ populations expressed only one of these markers, only the *Trh*-*Nfib* neurons were enriched for both (**Fig. 2E**), suggesting those to be the neuroendocrine population.

Lastly, no markers specific for neuroendocrine *Sst* ^+^ subpopulations have been described in literature. The identified clusters *Sst* -*Calb2* and *Sst* -*Sfrp2* do seem distinct in their expression of various genes. In *Sst* -*Sfrp2* neurons, *Sfrp2, Ar, Npy1r*, and *Mc4r* were enriched, while the *Sst* -*Calb2* neurons were enriched for *Calb1, Calb2, Kcnip4* and *Gpr101* (**Fig. S4B**).

### 2.3. Pre-autonomic neurons are embedded in the Adarb2^+^ clusters

To identify pre-autonomic neurons in our data, we integrated a recent single-cell dataset that utilized retrograde tracing from the IML [3]. From this dataset, the neurons identified by the authors to be from the PVN were integrated into our PVN atlas. The traced neurons were high in their expression of *Adarb2, Reln, Snca* and *Ntng1* (**Fig. 3A**). This was similar to the *Asb4* -*Adarb2, Brs3* -*Adarb2, Avp*-*Adarb2, Crh*-*Adarb2*, — *Slc17a6* -*Adarb2, Penk* -*Adarb2* and *Oxt* -*Adarb2* subpopulations, together to be referred to as the *Adarb2* ^+^ subpopulations. The data integration embedded the traced neurons within the *Adarb2* ^+^ populations, mostly the *Avp*-*Adarb2, Oxt* -*Adarb2, Penk* -*Adarb2* and *Crh*-*Adarb2* clusters (**Fig. 3B**). The traced neurons expressed all these neuropeptides to some extent, though the expression of *Oxt* was markedly the most pronounced (**Fig. 3C**).

**Figure 3:**
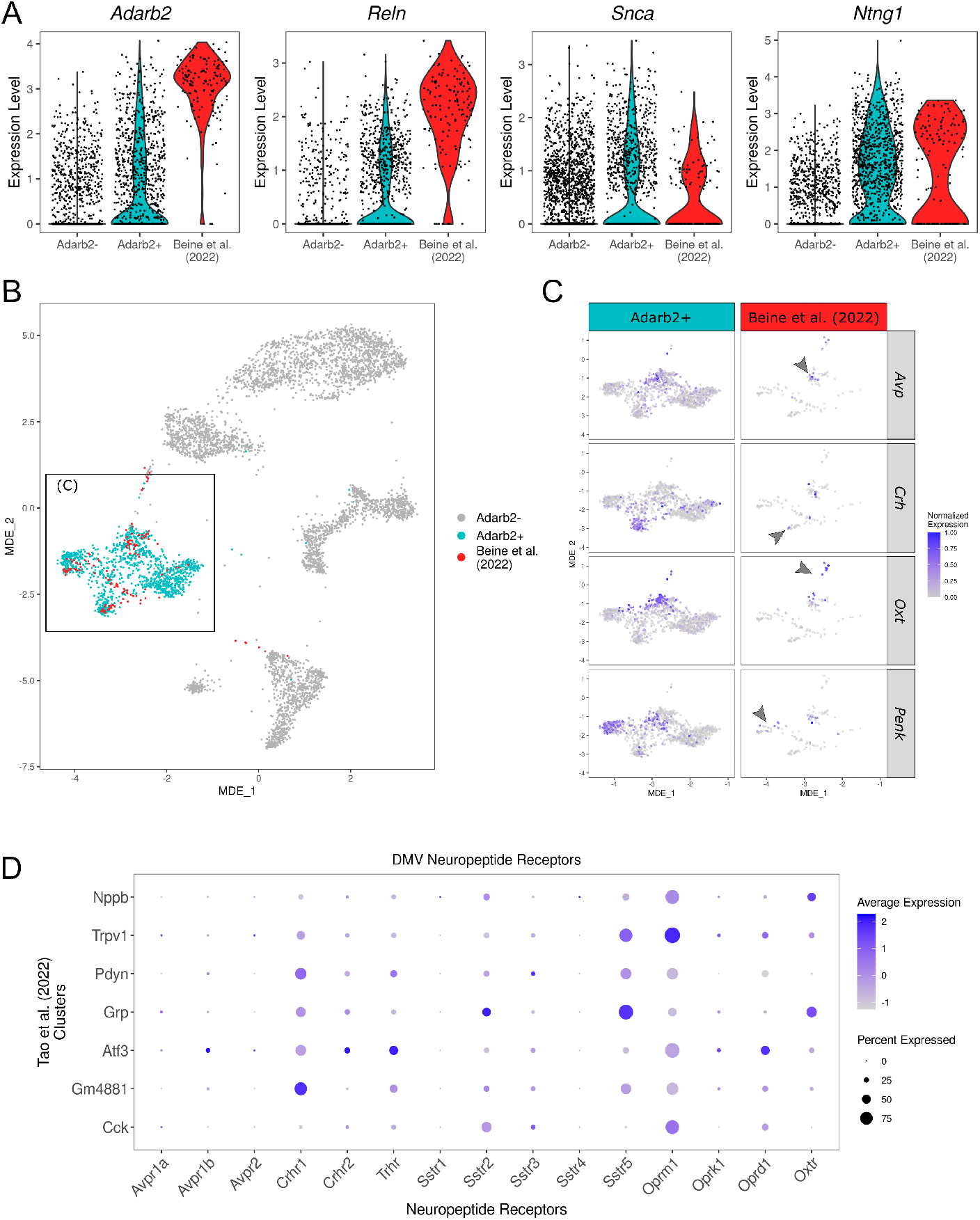
Pre-autonomic neurons co-cluster with the *Adarb2* ^+^ neurons. **A**: Violin plots showing expression of *Adarb2, Reln, Snca, Ntng1* in the PVN atlas and retrogradely traced PVN neuron from Beine *et al*. [3]. **B**: The MDE representation, with cells from Beine *et al*. [3] highlighted red. **C**: Zoomed-in MDE representation, showing the *Adarb2* ^+^ clusters. Rows show cells from Beine *et al*. [3] and the rest of the dataset, columns are split by expression of *Avp, Crh, Oxt*, and *Penk*. **D**: Dotplot showing neuropeptide receptor expression in the various populations of the DMV. Dot size represents percentage of cells expressing a gene, dot color represents the average expression level. Populations used same annotations as in Tao *et al*. [28].

Besides the IML, the pre-autonomic neurons of the PVN also project to the DMV. Although there are currently no retrograde tracing sequencing datasets available, a recent single-cell sequencing dataset of the general DMV [28] may still give insight into possibly relevant circuitry, through its neuropeptide receptor repertoire. The neuropeptidergic receptors that relate to PVN-expressed neuropeptides, and were found in the various DMV transcriptional populations are *Crhr1, Trhr, Sstr2, Sstr5, Oprm1, Oprd1* and *Oxtr* (**Fig. 3D**).

In summary, our results indicate that sympathetic pre-autonomic neurons include *Avp*-*Adarb2, Crh*-*Adarb2, Oxt* -*Adarb2*, and *Penk* -*Adarb2* neurons. Brainstem-projecting parasympathetic pre-autonomic neurons may be more diverse, since receptors for *Crh, Trh, Sst, Penk* and *Oxt* are present in the DMV.

## 3. Discussion

In this study, we integrated single-cell PVN datasets into a general hypothalamic atlas, and subsequently selected PVN-specific neurons from the integrated dataset. We characterized the resulting dataset, revealing multiple transcriptional subpopulations for neuropeptidergic neurons. For the *Avp*^+^, *Oxt* ^+^, *Crh*^+^ and *Trh*^+^ neurons, we identified distinct subpopulations that were enriched for neuroendocrine marker genes. Finally, we describe how spinally-projecting pre-autonomic neurons cluster entirely with various *Adarb2* ^+^ neurons, and we identify putative neuropeptidergic types of brainstem-projecting pre-autonomic neurons.

Since we used a microdissected high-depth sequencing dataset as benchmark [33], our final dataset is unlikely to contain any non-PVN neuron populations. Rare cell types for the PVN that are much more prominent in adjacent brain regions were excluded based on in-situ hybridization data. Nonetheless, this approach also has its limitations, as the sample size of the benchmark dataset was relatively small. Rare transcriptional populations, or populations restricted to undersampled PVN subregions are less likely to be uncovered through this method.

Another limitation of our study is linked to the use of the retrogradely traced single-cell data from Beine *et al*. [3]. Their study injected the retrograde tracer into the L1 spinal cord, while the IML spans from T2 through L2 in mice [24]. Thus, the traced population is likely only a limited representation of the spinally-projecting pre-autonomic neurons, as the axon terminal location could be a source of transcriptional variation in pre-autonomic neurons.

Despite the limitations, we were able to identify multiple subpopulations for all neuropeptidergic neuron populations previously shown to reside in the PVN. For *Avp*^+^ neurons, we identified three distinct subpopulations that have not been described before: *Avp*-*Adarb2, Avp*-*Tac1* and *Avp*-*Th*. While the presence of *Th* in *Avp*^+^ neurons is not novel [1], the presence of *Th* as marker for a distinct subpopulation of *Avp*^+^ neurons has not been described before. The *Avp*-*Tac1* and *Avp*-*Th* subpopulations were both highly correlated with the magnocellular neuroendocrine *Oxt* ^+^ neurons, and both were high in expression of the neuroendocrine markers as described by Lewis *et al*. [16]. These results indicate that these *Avp*^+^ populations may be magnocellular neuroendocrine neurons. On the other hand, the *Avp*-*Adarb2* population was similar to the retrograde traced neuron, indicating a function as pre-autonomic neurons.

The identified *Oxt* ^+^ neuron subpopulations were transcriptionally the same as identified by Lewis *et al*. [16] – with a side note that the magnocellular population in their dataset also contained some neuroendocrine *Avp*^+^ neurons. We identify the *Oxt* -*Foxp1* neurons to be the magnocellular neuroendocrine *Oxt* ^+^ population, and identify the parvocellular *Oxt* -*Adarb2* neurons to be pre-autonomic, based on the transcriptional overlap with retrogradely traced neurons. However, the *Oxt* -*Adarb2* population is functionally more diverse than just pre-autonomic function, as neurons from this population have also been shown to project to the nucleus accumbens to mediate social reward learning [16].

For the *Crh*^+^ neurons, two subpopulations were identified: *Crh*-*Nr3c1* and *Crh*-*Adarb2*. The *Crh*^+^ subpopulations are likely similar to the population as discussed by Romanov and Harkany [22], albeit annotated with different secondary cluster markers. We find that the *Crh*-*Nr3c1* neurons are enriched for multiple neuroendocrine markers genes, while the *Crh*-*Adarb2* neurons are transcriptionally similar to a subset of spinally-projecting pre-autonomic neurons. In contrast, Romanov and Harkany [22] posit that both subpopulations are neuroendocrine, and that other markers – like *Npy1r* – define subsets of these populations that are pre-autonomic instead of neuroendocrine. While this explanation is possible, it seems unlikely for two reasons. First, neuroendocrine markers like *Agtr1a, Nr3c1*, and *Scgn* are lacking entirely in the *Crh*-*Adarb2* neurons, while *Avp* was also relatively low in expression in these cells. Secondly, no retrogradely traced pre-autonomic neurons were found co-clustering with the *Crh*-*Nr3c1* neurons. This suggest that these two *Crh*^+^ subpopulations are distinct, not only transcriptionally but also functionally.

Multiple subpopulations of *Trh*^+^ neurons were also identified in our analysis, but the functional interpretation was more difficult than for other populations, due to lack of literature on these subpopulations. We observed an enrichment of two neuroendocrine markers in *Trh*-*Nfib* neurons, but the other *Trh*^+^ sub-populations were not functionally classifiable. The *Brs3* -*Adarb2* population, which expresses a low level of *Trh*, is transcriptionally similar to other *Adarb2* ^+^ populations, and a pre-autonomic function seems plausible. Nevertheless, since no retrogradely traced neurons seemed to cluster with this subpopulation, no conclusive evidence was found in our analysis to substantiate this hypothesis.

Similarly, the identified *Sst* ^+^ subpopulations could not be linked to a functional class due to lacking literature on functional *Sst* ^+^ subpopulations. Despite this, two of the identified differential genes do stand out. First, the *Ar* has been described in literature to define a subgroup of *Sst* ^+^ neurons that is present in both sexes, but in larger numbers in males [13]. This population is postulated to be relevant for sexually dimorphic action of SST on the HPS axis. The other notable gene is *Gpr101*, which has been reported as the causative gene to X-linked acrogigantism – a disorder involving the growth hormone axis [15]. Since both populations can be linked to the HPS axis, through either *Ar* or *Gpr101*, we hypothesize both subpopulations are neuroendocrine in nature, with the two populations acting as separate substrates for HPS axis inhibition.

Finally, we analyzed the neuropeptide receptor repertoire of the DMV. Receptors for CRH, OXT, PENK and SST were present, as well as the receptor for TRH to a lesser extent. This receptor expression profile is consistent with literature on PVN-DMV neurons, which reports expression of AVP, OXT, PENK, SST, and CRH [23, 5]. The presence of TRH receptor might indicate relevance for this neuropeptide in PVN-DMV circuitry, though TRH projections could originate from brain regions other than the PVN as well. As the DMV-projecting AVP and OXT neurons were reportedly parvocellular, we can infer that the *Avp*-*Adarb2* and *Oxt* -*Adarb2* populations are the most likely candidates for the vasopressinergic and oxytocinergic PVN-DMV connections. This would implicate these populations in both pre-sympathetic and pre-parasympathetic function, with currently no known differentiating factor to define these functional subpopulations. In summary, our findings reinforce and expand on current literature on PVN-DMV circuitry and introduces novel avenues to elucidate neuropeptidergic subpopulations involved in this circuit.

Concluding, our study presents a detailed overview of the neuronal transcriptional subpopulations in the PVN, and attempts to resolve functional identities for the identified populations (**Fig. 4**). The study pinpoints interesting markers of neuropeptidergic subpopulations, that may be used for research into functionalities that are specifically associated with these subpopulations.

**Figure 4:**
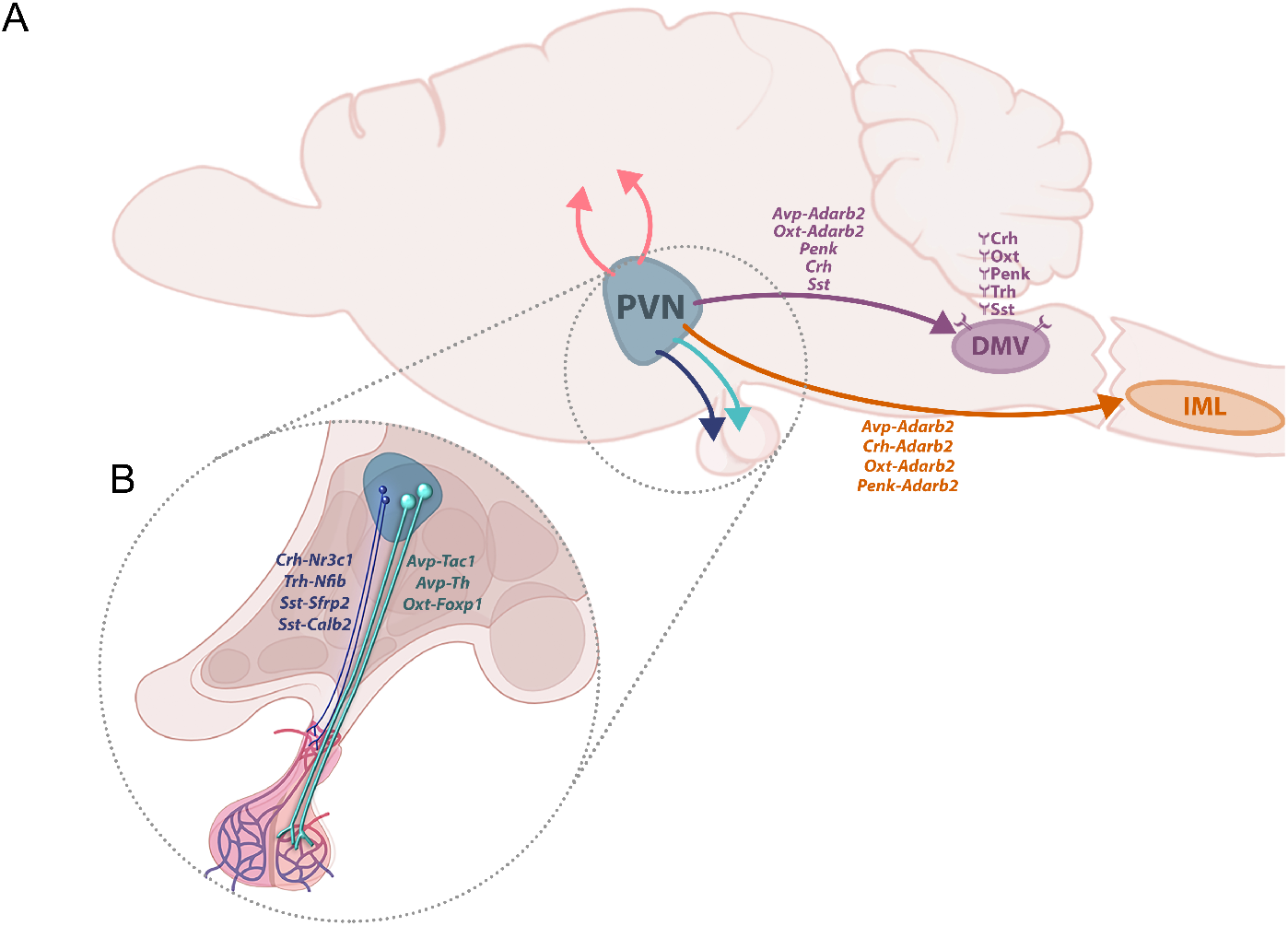
Summarized findings from our PVN atlas. **A**: Overview of projections from the PVN to the IML and DMV, with relevant cell populations, neuropeptides, and neuropeptide receptors shown. **B**: Overview of projections from the PVN to the pituitary, with relevant cell populations shown.

## 4. Methods

### 4.1. scRNA-seq data collection and preprocessing

PVN scRNA-seq datasets were acquired from the relevant Gene Expression Omnibus (GEO) repositories (**Table 1**). The hypothalamic atlas data (HypoMap) was acquired from CELLxGENE (**Table 1**). Each dataset was preprocessed separately to filter low quality cells based on transcript count, gene count, and percentage mitochondrial gene expression (**Table S1**). Datasets using Ensembl IDs were converted to MGI symbols. Furthermore, dataset metadata was standardized to include `assay, disease, organism, sex, SRA ID, Sample_ID, and Dataset`, using the HypoMap atlas metadata as template. Within the atlas, cells were annotated at various levels of granularity, ranging from `C7_named to C465_named`. The number in these annotation levels denotes the number of clusters at that annotation level. Non-atlas datasets were annotated at the `C7_named` level by comparison of cluster marker genes with the respective expression patterns in the atlas, assigning clusters in the non-atlas dataset based on these comparisons. Lastly, the datasets were merged on the raw counts and metadata. For further analysis, we selected neuronal cells only, based on the `C7_named` annotation key. Additionally, the first principal component of the immediate early genes (IEGs) *Fos, Fosb, Jun, Junb, Egr1, Egr2, Npas4, Nr4a1, Nr4a2, Nr4a3* was calculated on normalized gene expression, to regress out the effect of IEGs during integration.

**Table 1:** Datasets used in this paper. Abbreviations: CxG = CELLxGENE; NS = not specified;

| Reference | GEO/CxG | Technology | Cell type / Region | Cells included | Genes / cell |
| --- | --- | --- | --- | --- | --- |
| Beine *et al.* [3] | GSE212409 | 10X | Includes PVN | 142 | 2546 |
| Lewis *et al.* [16] | GSE147092 | SMART-Seq2 | *Oxt*<sup>+</sup> | 134 | 1146706 |
| Lopez *et al.* [17] | GSE161751 | 10X | PVN | 6399 | 8860 |
| Short *et al.* [25] | GSE174085 | SMART-Seq2 | *Crh*<sup>+</sup> | 519 | 1494713 |
| Steuernagel *et al.* [27] | HypoMap | 10X, Drop-Seq | Hypothalamus | 384925 | 4812 |
| Son *et al.* [26] | GSE166616 | Drop-seq | *Sim1*<sup>+</sup> | 2440 | 7679 |
| Tao *et al.* [28] | GSE172411 | NS | DMV | 282 | 1539569 |
| Xu *et al.* [33] | GSE148568 | SMART-SCRB | PVN | 706 | 177416 |

### 4.2. scRNA-seq data integration

The highly variable genes of the merged dataset were selected using the function `highly_variable_genes` from scanpy [31] with parameters `n_top_genes = 2500, flavor = ‘seurat’, subset = True, batch key = ‘Dataset’`. Using SCVI [18], the dataset was then integrated on Sample ID with assay, `suspension_type` and sex as categorical covariates, and with `immediate_early` and `pctMito_RNA` as continuous covariates. The model used parameters `n_layers = 3, n_latent = 20, dropout_rate = 0.01, dispersion = ‘gene-cell’` and `gene_likelihood = ‘zinb’`. The model was trained for 400 epochs, with n_epochs_kl_warmup = 400, lr = 0.01, lr_min = 0.0001, reduce_lr_on_plateau = True, lr patience = 8, lr factor *≈* 0.56, lr_scheduler metric = ‘elbo validation’. The resulting trained SCVI model was passed to SCANVI [32]. The SCANVI model was then trained for 250 epochs, using `C66_named` as label key. The SCANVI model was trained using the same parameters as SCVI, except for `classification_ratio = 200` and `n_epochs_kl_warmup = 100`. Both SCVI and SCANVI models were trained with early stopping, using `early_stopping = True, early_stopping_patience = 25, early_stopping_monitor = ‘elbo_validation’` as parameters.

To isolate PVN-specific cells from the integrated dataset, the cells from Xu *et al*. [33] were used as benchmark. In that study, PVN boundaries were identified based on fluorescent agouti-related peptide (AgRP) axon tracts, and PVN neurons were microdissected and sequenced using a plate-based sequencing method (SMART-SCRB). The resulting dataset was thus deemed the most accurate representation of the PVN, due to the combination of high sequencing depth and precise cell isolation method used. The other PVN datasets were either selective for a single neuropeptide neuron type, or not fully specific for the PVN, due to contamination with neurons from bordering regions.

As such, a subset was taken of predicted `C66_named` clusters that contained 5 or more cells originating from Xu *et al*. [33]. This subset was clustered using the Leiden algorithm [29] with `n_neighbors = 30` and `resolution = 1`. A subset was taken of the resulting clusters, again retaining only clusters with at least 5 cells from Xu *et al*. [33]. These subsetting steps retained 98.58% and 97.56% of the cells from Xu *et al*. [33], respectively. The resulting subset was then reintegrated using the same steps and parameters as before, except for the highly variable gene selection with `n_top_genes = 1000`.

The SCANVI latent embedding was used to cluster the data with the Leiden algorithm [29] with `n_neighbors = 30` and `resolution = 1`. Each of the resultant 20 clusters were analyzed for presence of Xu *et al*. [33] neurons. Only clusters with presence of minimally 5 of these neurons were annotated. An exception was made for *Trh*-*Ucn3*, as Allen Brain Atlas in-situ hybridization data showed *Ucn3* expression in the PVN (**Fig. S5**). For annotation, these clusters were tested for potential subclusters using the `FindSubClusters` function, and subclusters containing no cells from Xu *et al*. [33] were excluded. The resultant clusters and subclusters were tested for enriched genes using the `FindMarkers` functions, and were annotated accordingly. Finally, a subset was taken with all annotated clusters, containing only the annotated populations described in the results section.

To analyze the retrogradely traced neurons from Beine *et al*. [3], the PVN neurons within that dataset were selected based on *Sim1* expression. The selected neurons were integrated into the PVN atlas, using the same steps and parameters for SCANVI integration as described before. Altered parameters were `n_top_genes = 1000` and `early_stopping = False`. The latent representation of this final integration was used for visualization of the data, using the `preserve_neighbors` function from the minimally distorting embedding (MDE) framework [2]. Parameters used for MDE visualization were `attractive penalty = Huber, repulsive penalty = Log, n_neighbors = 30`, and `device = ‘cuda’`.

Lastly, for the analysis of the neuropeptide receptors in the DMV, preprocessing and clustering steps as described by Tao *et al*. [28] were replicated.

### 4.3. Statistics

Single-cell transcriptomic pipelines and statistical computations were performed using both Python 3.7.10 and R 4.1.2 [20].

## 5. Ethics & Integrity

### 5.1. Data & Code availability

Reproducible code for the generation of the dataset and figures is available at https://github.com/jberkh/2023_PVN_Atlas. The integrated PVN atlas can be found at Zenodo (https://doi.org/10.5281/zenodo.8160038) and will be published to CELLxGENE.

### 5.2. Funding

This research was funded by ZonMw Open Competition grant #09120012010051.

### 5.3. Conflict of interest

The authors declare no conflict of interest.

## Supplement

**[Figure S1:].**
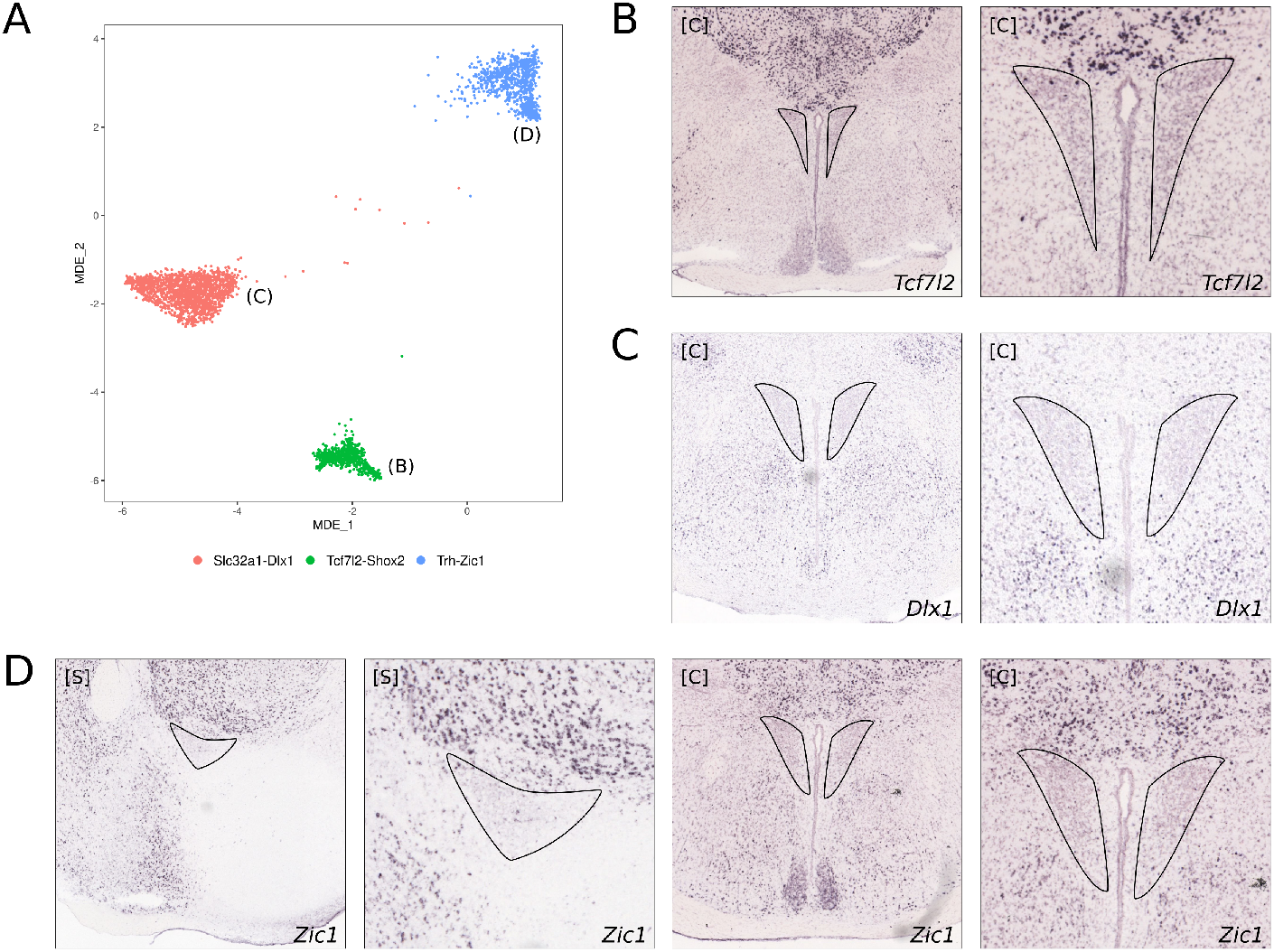
Neurons from the *Tcf7l2* -*Shox2, Slc32a1* -*Dlx1* and *Trh* -*Zic1* are scarcely present in the PVN. **A**: Minimally distorting embedding (MDE) representations of the excluded clusters, *Tcf7l2* -*Shox2, Slc32a1* -*Dlx1* and *Trh*-*Zic1*. **B**: Allen Brain Atlas in-situ hybridization data of *Tcf7l2* in coronal sections. Approximate boundaries of the PVN shown as black line. **C**: Allen Brain Atlas in-situ hybridization data of *Dlx1* in coronal sections. Approximate boundaries of the PVN shown as black line. **D**: Allen Brain Atlas in-situ hybridization data of *Dlx1* in sagittal and coronal sections. Approximate boundaries of the PVN shown as black line.

**[Figure S2:].**
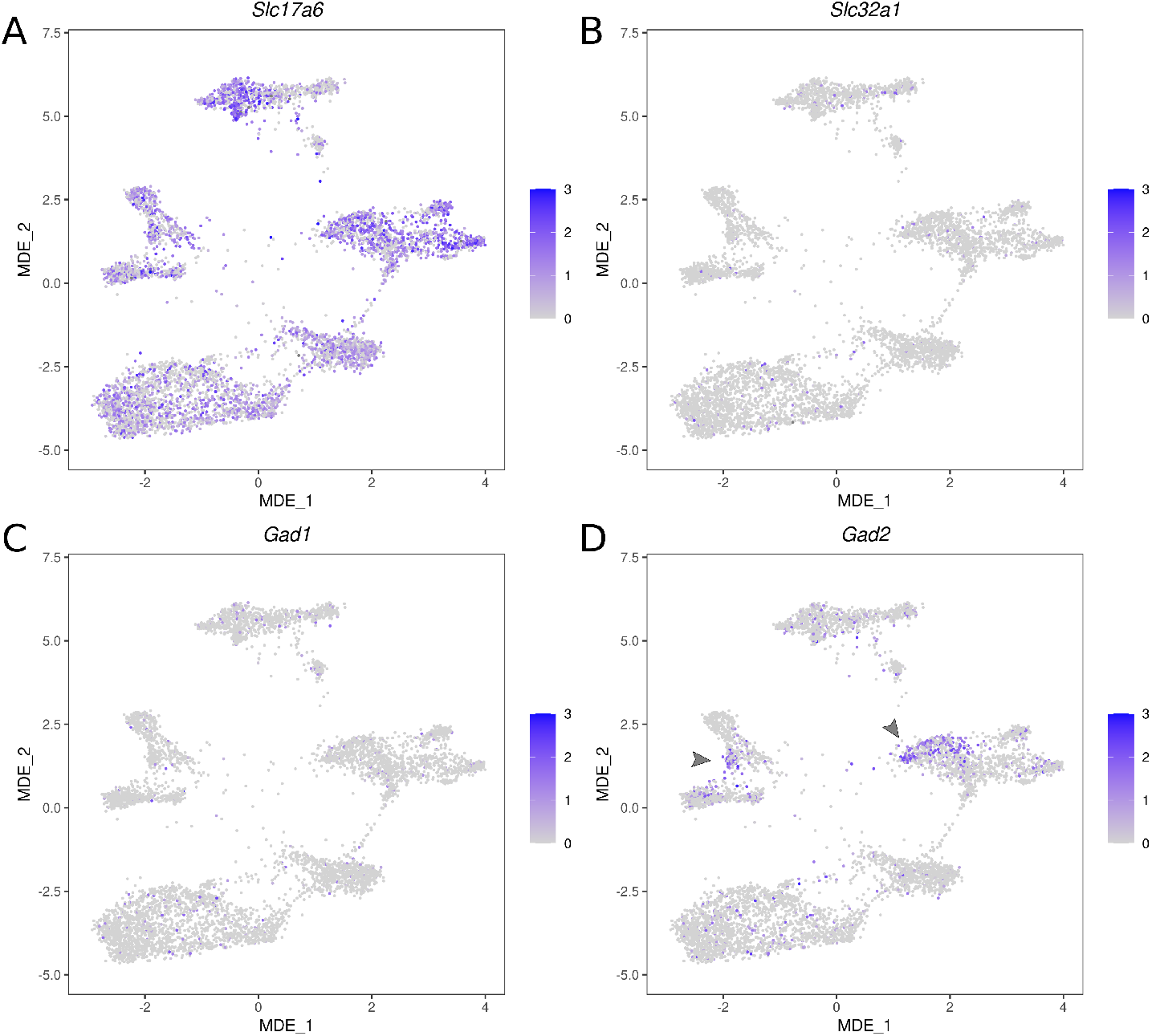
All PVN clusters are *Slc17a6* ^+^/*Slc32a1* ^*−*^. **Despite lacking *Slc32a1*, some PVN clusters express GABA biosynthesis enzymes, like *Gad2***. **A**: MDE representation of the PVN atlas showing expression of *Slc17a6* **B**: MDE representation of the PVN atlas showing expression of *Slc32a1* **C**: MDE representation of the PVN atlas showing expression of *Gad1* **D**: MDE representation of the PVN atlas showing expression of *Gad2*

**[Figure S3:].**
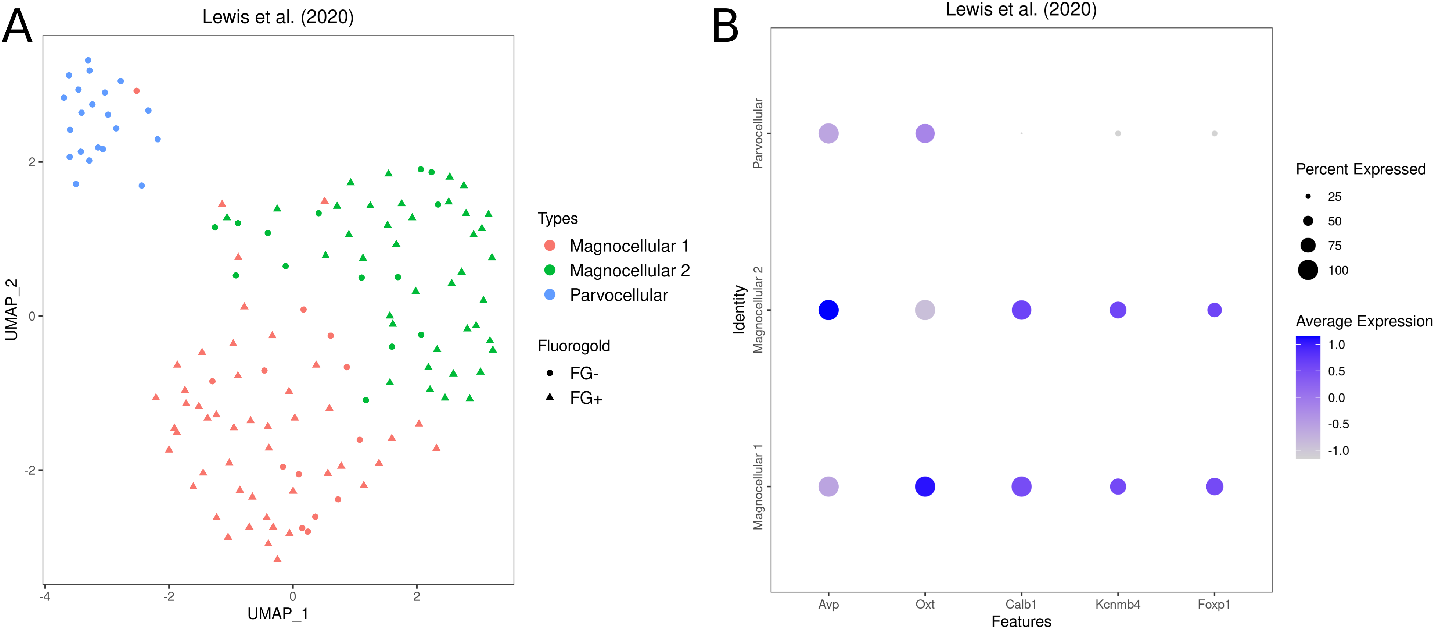
The magnocellular neurons from Lewis *et al*. [16] are both vasopressinergic and oxytocinergic. **A**: UMAP representation of the dataset, using the original annotations from Lewis *et al*. [16]. Magnocellular cells are divided into two subpopulations. Fluorogold positivity, indicative of neuroendocrine function, depicted by shape. **B**: Dot plot showing expression levels of *Avp, Oxt, Calb1, Kcnmb4* and *Foxp1* in the three subpopulations from panel A.

**[Figure S4:].**
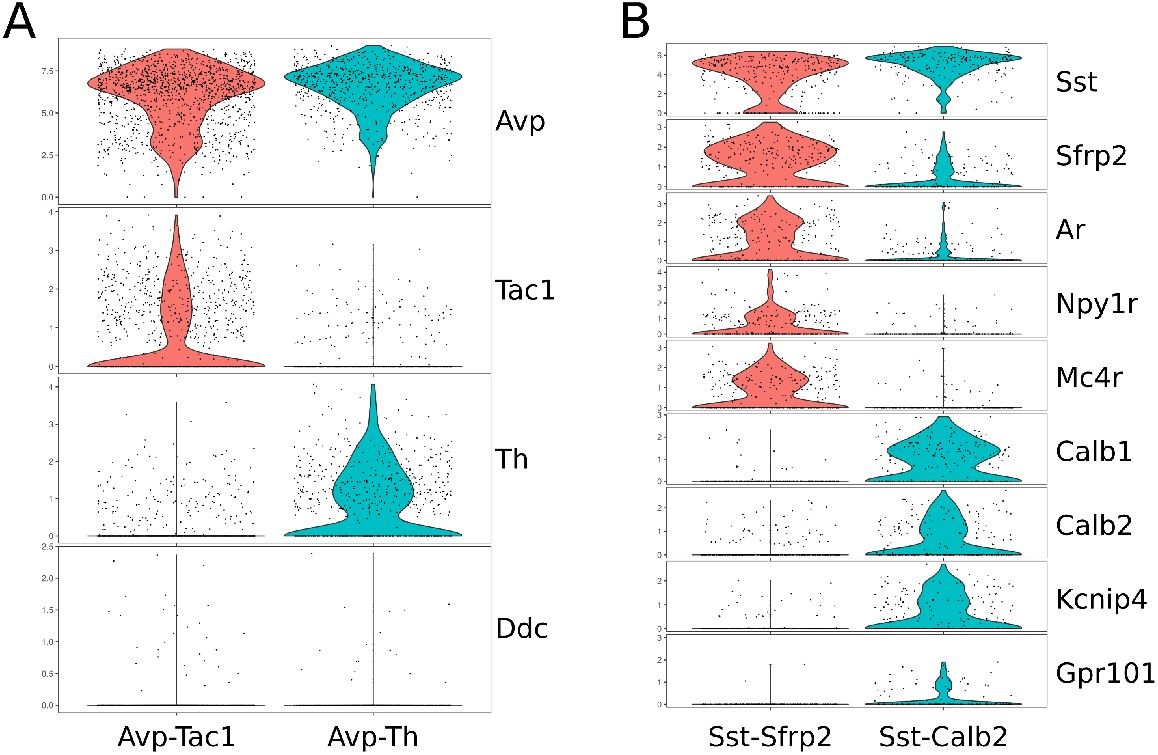
Differential genes between subpopulations of *Avp*^+^ and *Sst* ^+^ neurons. **A**: Violin plots showing the expression of *Avp, Tac1, Th* and *Ddc* in the *Avp*-*Tac1* and *Avp*-*Th* populations. **B**: Violin plots showing the expression of *Sst, Sfrp2, Ar, Npy1r, Mc4r, Calb1, Calb2, Kcnip4*, and *Gpr101* in the *Sst* -*Sfrp2* and *Sst* -*Calb2* populations.

**[Figure S5:].**
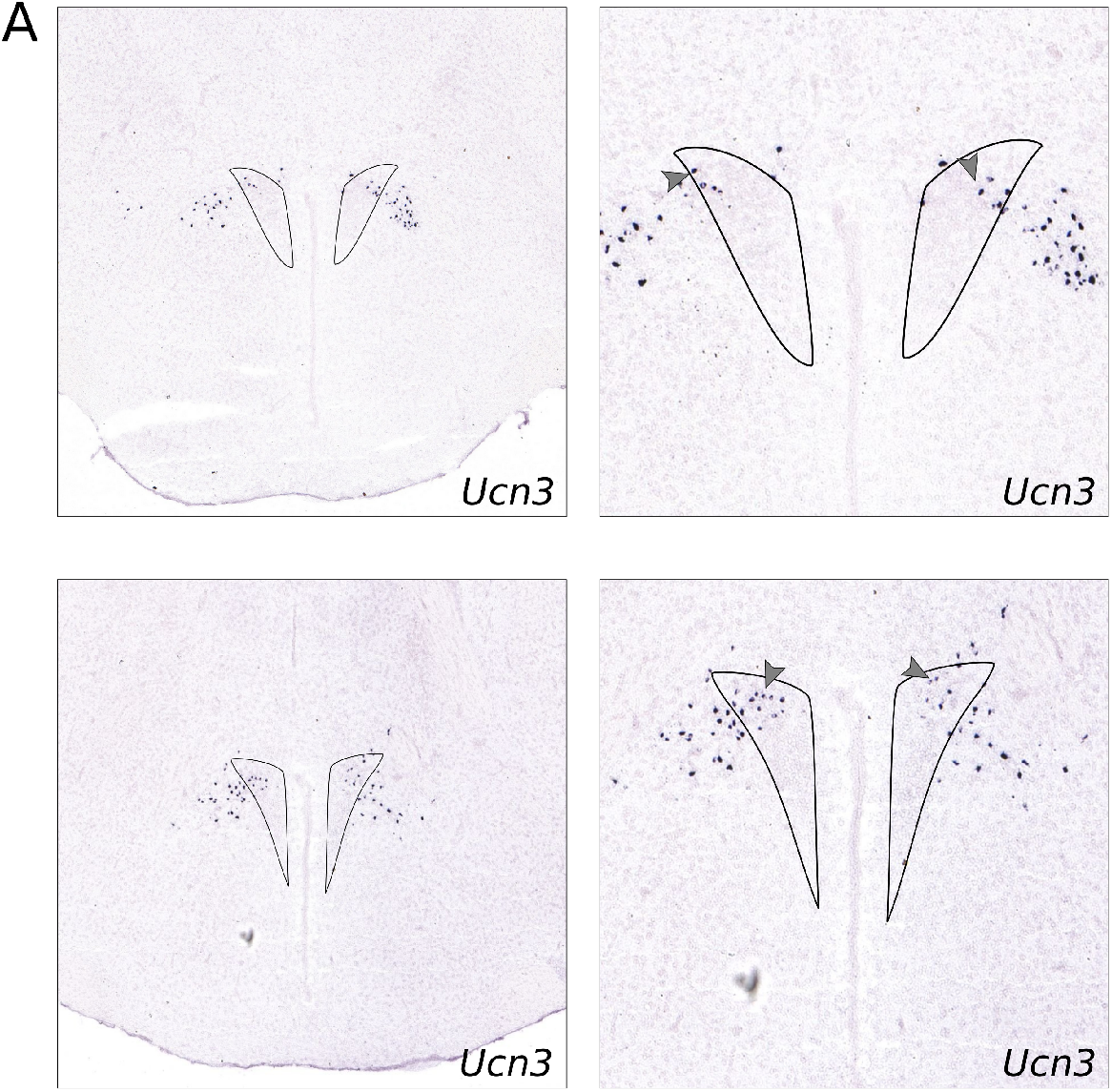
Expression of *Ucn3* is found in the both the perifornical area and PVN. **B**: Allen Brain Atlas in-situ hybridization data of *Ucn3* in coronal sections. Approximate boundaries of the PVN shown as black line.

**[Table S1:].**
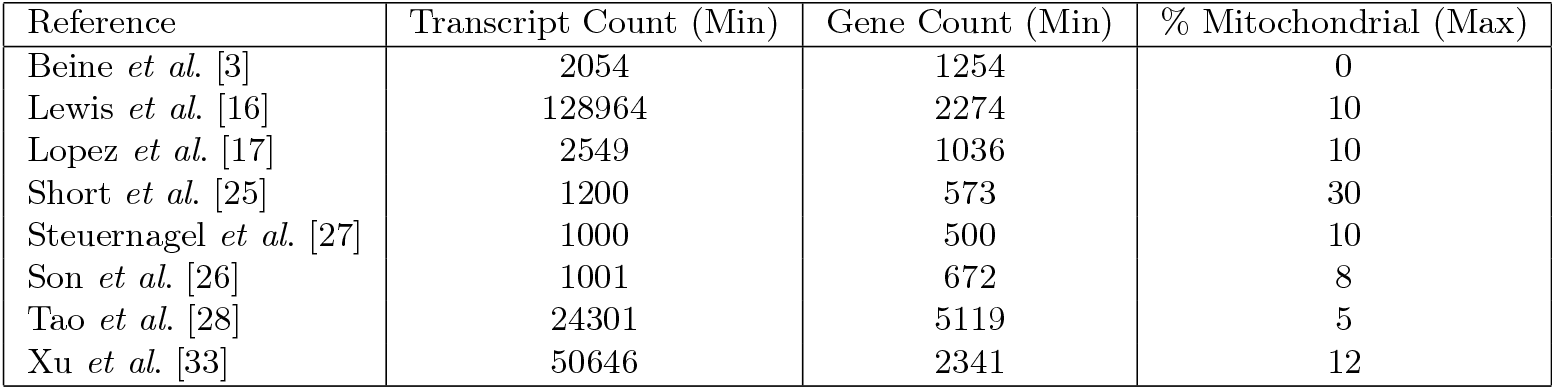
Dataset thresholded metadata after preprocessing.

| Reference | Transcript Count (Min) | Gene Count (Min) | % Mitochondrial (Max) |
| --- | --- | --- | --- |
| Beine *et al.* [3] | 2054 | 1254 | 0 |
| Lewis *et al.* [16] | 128964 | 2274 | 10 |
| Lopez *et al.* [17] | 2549 | 1036 | 10 |
| Short *et al.* [25] | 1200 | 573 | 30 |
| Steuernagel *et al.* [27] | 1000 | 500 | 10 |
| Son *et al.* [26] | 1001 | 672 | 8 |
| Tao *et al.* [28] | 24301 | 5119 | 5 |
| Xu *et al.* [33] | 50646 | 2341 | 12 |

## Notes

https://github.com/jberkh/2023_PVN_Atlas

https://doi.org/10.5281/zenodo.8160038

### Competing Interest Statement

The authors have declared no competing interest.

